# Antibiotics-induced alterations in gut microbiol composition leads to blood-brain barrier permeability increase in rhesus monkeys

**DOI:** 10.1101/2020.01.02.890939

**Authors:** Qiong Wu, Yingqian Zhang, Yinbing Zhang, Chunchao Xia, Qi Lai, Zaiquan Dong, Weihong Kuang, Cheng Yang, Dan Su, Hongxia Li, Zhihui Zhong

## Abstract

Blood-brain barrier (BBB) contributes to maintenance of brain homeostasis. Gut microbiome composition affected BBB development and expression of tight junction proteins in rodents. However, we still do not know if there’s any direct effect of gut microbiol composition on BBB permeability and function in normal adult animals. In current study, we determined temporal and spatial change of BBB permeability in rhesus monkeys receiving either oral or intravenous amoxicillin-clavulanic acid (AC), by monitoring CSF/serum albumin ratio (AR) and the volume transfer constant (K_trans_). We showed that oral, but not intravenous AC led to a significant alteration in gut microbiol composition and increase of BBB permeability in all monkeys, especially in thalamus area. Acetic acid and propionic acid might play a pivotal role in this newly found communication between gut and central nervous system. Antibiotics-induced gut microbiol composition change, especially the decreasing of acetic and propionic acid producing phyla and genera, leads to increase of BBB permeability, which may contribute to a variety of neurological and psychological diseases.

## INTRODUCTION

Gut-brain axis, or microbiome-gut-brain axis, often refers to the sophisticated mutual interactions between gut microbiome and the central nervous system (CNS) ^1-3^, which has been demonstrated in preclinical and clinical studies^4^. Changes in gut microbiota could influence gastrointestinal physiology as well as homeostasis of CNS ^3^, although the exact mechanism has not yet been fully understood^5^.

Blood-brain barrier (BBB) serves as the main gatekeeper of the brain, regulating the passage of nutrients and regulatory molecules from the circulatory system and guarding the CNS from toxins and pathogens^6^. Weakened BBB integrity, or ‘leaky BBB’, significantly contributes to the progression of many neurodegenerative and psychological diseases, including stroke, traumatic brain injury, multiple sclerosis, Alzheimer’s disease (AD), Parkinson’s disease (PD), schizophrenia and autism spectrum disorder (ASD)^7-9^. However, mechanisms by which the BBB opens up differ in different pathophysiological conditions and remains elusive for the most part^10^.

A crucial role of gut microbiome and their metabolites in the formation of the BBB during embryonic and neonatal stage has been demonstrated in germ-free (GF) mice, in which the BBB is more permeable to macromolecules compared to conventionally raised animals^11^. This result, however, is contradictory to Leclercq et al, showing low-dose oral penicillin in early life of mice moderately upregulated proteins which may enhance the integrity of BBB, in hippocampus^12^. So far, no direct evidence of BBB disruption was ever documented. Effects of gut microbiol change on BBB permeability and thus CNS, and by what mechanisms these effects happen still remain inconclusive.

It has been established that antibiotic administration affects not only the pathogens to which they are directed but also the entire microbiota^13^. In current study, a clinical dose of amoxicillin-clavulanic acid (AC), which are the most commonly prescribed antibiotic combination to patients, was used either orally to induce gut microbiol change^14^ or intravenously. A well-established longitudinally monitoring method for monkey BBB permeability^9^ was adopted to monitor temporal and spatial changes of BBB permeability after AC administration, thus to explore effect of antibiotic-induced gut “dysbiosis” comparing to “homeostatic gut microbiol condition” on BBB permeability.

## MATERIALS AND METHODS

### Experimental Animals

Twenty-one adult, male rhesus monkeys, weighing 4-10 kg and aged 4-6 years, were provided by Sichuan Green-House Biotech Co., Ltd, Sichuan Province, China, 2 weeks prior to experiment, and were housed in separate cages and maintained in a 12-hour light/dark cycle at room temperature. All the experimental procedures were conducted in accordance with the Guidance Suggestions for the Care and Use of Laboratory Animals, formulated by the Ministry of Science and Technology of People’s Republic of China, as well as AAALAC-related animal ethical criteria. The protocols were approved by the Animal Care Committee of the Sichuan University West China Hospital (Chengdu, China). None of the animals were sacrificed or severely wounded or during the experiment. All of them were sent back to original breeding base at the end of the study.

### Antibiotic administration and sample collection

A total of 21 adult monkeys were randomized into 2 groups (Table 1). In oral AC group (n = 15), AC diluted in 10 ml of 5% dextrose solution were given orally twice a day for 14 days at 200 mg/28.5 mg/kg body weight, a dose commonly used in clinical practice. In intravenous AC group (n = 6), AC was given intravenously twice a day for 14 days at 6 mg/0.86 mg/kg body weight, leading to the C_max_ of amoxicillin in blood similar to the orally administered AC. Fecal, cerebrospinal fluid (CSF), and blood samples were obtained before and after antibiotic exposure. All samples were stored at - 80□ until further analysis.

**Table 1.**
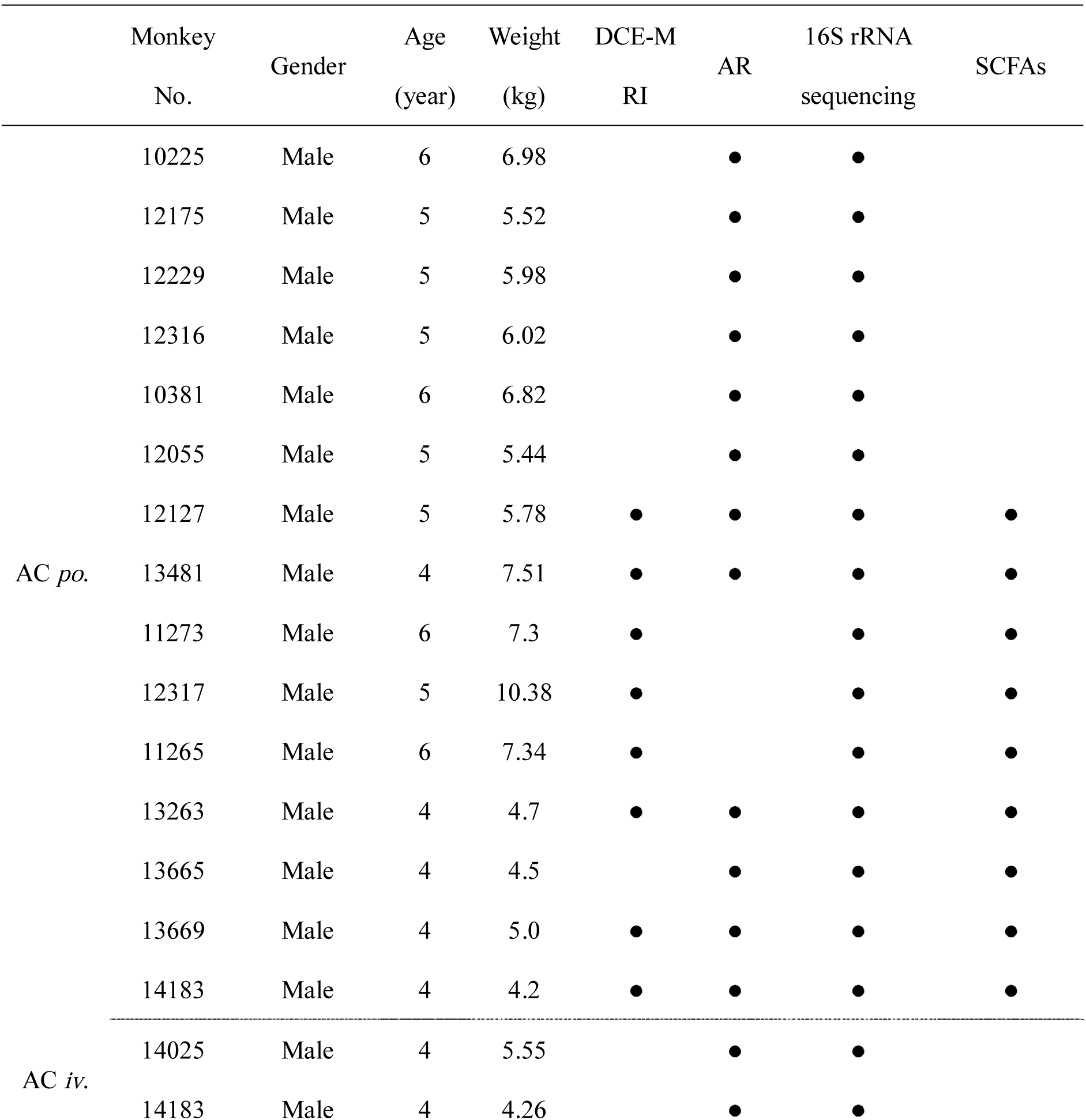
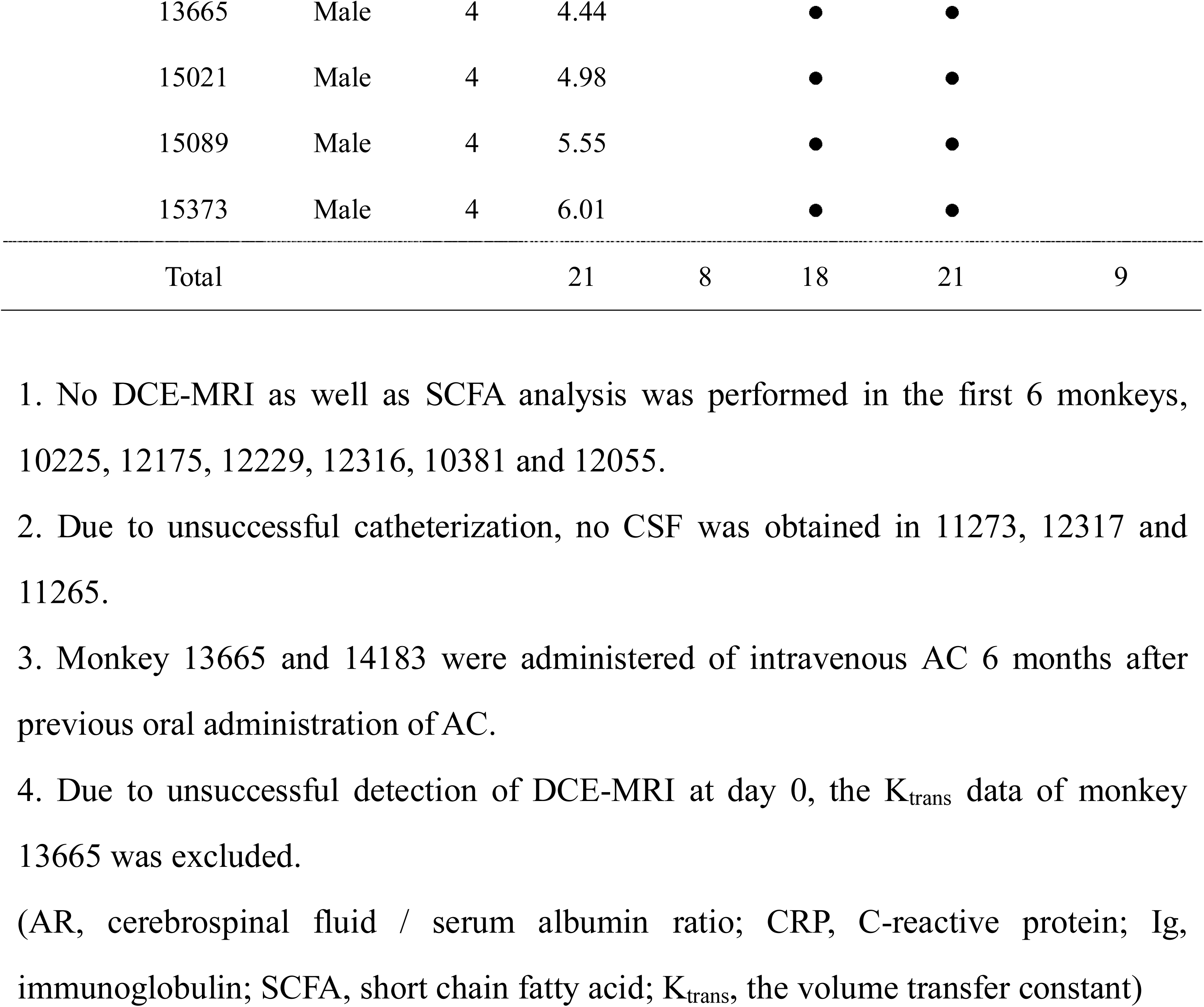
The basic information of rhesus monkeys.

### Serial CSF Sampling from Cisterna Magna

Minimally invasive catheterization to sample the CSF from monkey cisterna magna was performed as previously described^9^. For detailed information, see Supplementary material.

### The CSF/serum albumin ratio (AR) measurement

The CSF and serum albumin were measured by corresponding enzyme-linked immunosorbent assay (ELISA) (ab108788, abcam, USA) and colorimetric method respectively. AR = albumin in CSF (mg/L) / albumin in serum (g/L). The quality of CSF was assessed as described previously^9^. Considering the variance of normal level of AR among the animals, fold increase of AR was calculated to accurately indicate the permeability change of BBB.

### DCE-MRI

Monkeys received dynamic contrast-enhanced scanning with 3.0T MRI (DCE-MRI, Magnetom Verio, Siemens Medical Solution, Germany) to further identify the pattern, especially the spatial distribution, of BBB permeability change at day 0, 4, 7, 14 and 21. Frontal cortex, white matter, caudate nucleus, thalamus, and hippocampus were selected as regions of interest (ROIs). Raw data were processed by Siemens Tissue 4D software. Extended Tofts model was chosen to calculate the K_trans_ value (i.e., the volume transfer constant, which represents flow from the intravascular to the extravascular space). Fold increase of K_trans_ was used to accurately represent the permeability change of BBB due to the variance of normal level of K_trans_.

### 16S rRNA gene sequence analysis

The 16S rRNA analyses were used to characterize the gut microbiome in each fecal sample. Microbial DNA was extracted from stool samples using the Fast DNA^®^ SPIN Kit for Soil (MP Biomedicals, USA) according to manufacturer’s protocols (see supplementary methods for detail).

### Gas chromatography analysis of short chain fatty acids (SCFAs)

Three types of SCFAs (acetic acid, propionic acid, and butyrate) in monkey fecal samples were quantified by gas chromatography (see supplementary methods for detail).

### Statistical Analysis

All data were expressed as mean ± standard error of mean (SEM). All graphs were generated using GraphPad Prism software (version 7.0, San Diego, CA, USA). The difference among the means of the groups was determined with the one-way ANOVA, followed by the Tukey test. Principal coordinates analysis (PCoA) of β-diversity analysis was performed to visualize the differences among gut microbiome. Comparisons between values of each time point were performed with Student’s t-test or Mann Whitney U test depending on normality of distribution. All correlations were analyzed statistically by using Pearson’s correlation. A p value of less than 0.05 was considered significant in all analysis.

## RESULTS

### Oral antibiotics led to reversible alterations of gut microbial composition

As shown in figure 1A, the trend of Rank-abundance distribution curve representing evenness and richness of bacterial community displayed significant differences between *po.* group and *iv.* group. In *po.* group, the richness and evenness of bacterial diversity decreased significantly during AC treatment and reached its trough 7 days after antibiotic administration (day 7). After cessation of AC, there was a general tendency towards the restoration to the original (day 0) microbiome composition. In contrast, intravenously treated monkeys barely showed variations of gut microbiome composition (Fig. 1A inserted). After oral AC medication, the α-diversity Shannon index decreased significantly from 4.14 ± 0.15 (day 0) to 1.52 ± 0.38 (day 7) (p < 0.001), indicating that there were fewer heterogeneously distributed bacterial families during the initial dosing period. Shannon index then began to increase slowly during the second half of AC medication period and the upward trend was more obvious after cessation of the AC treatment (Fig. 1B). However, the Shannon index was still significantly lower than that of day 0 (p = 0.010). Conversely, Shannon index remained stable in *iv.* group and no statistical differences were found among each time point (Fig. 1B).

**Fig.1.**
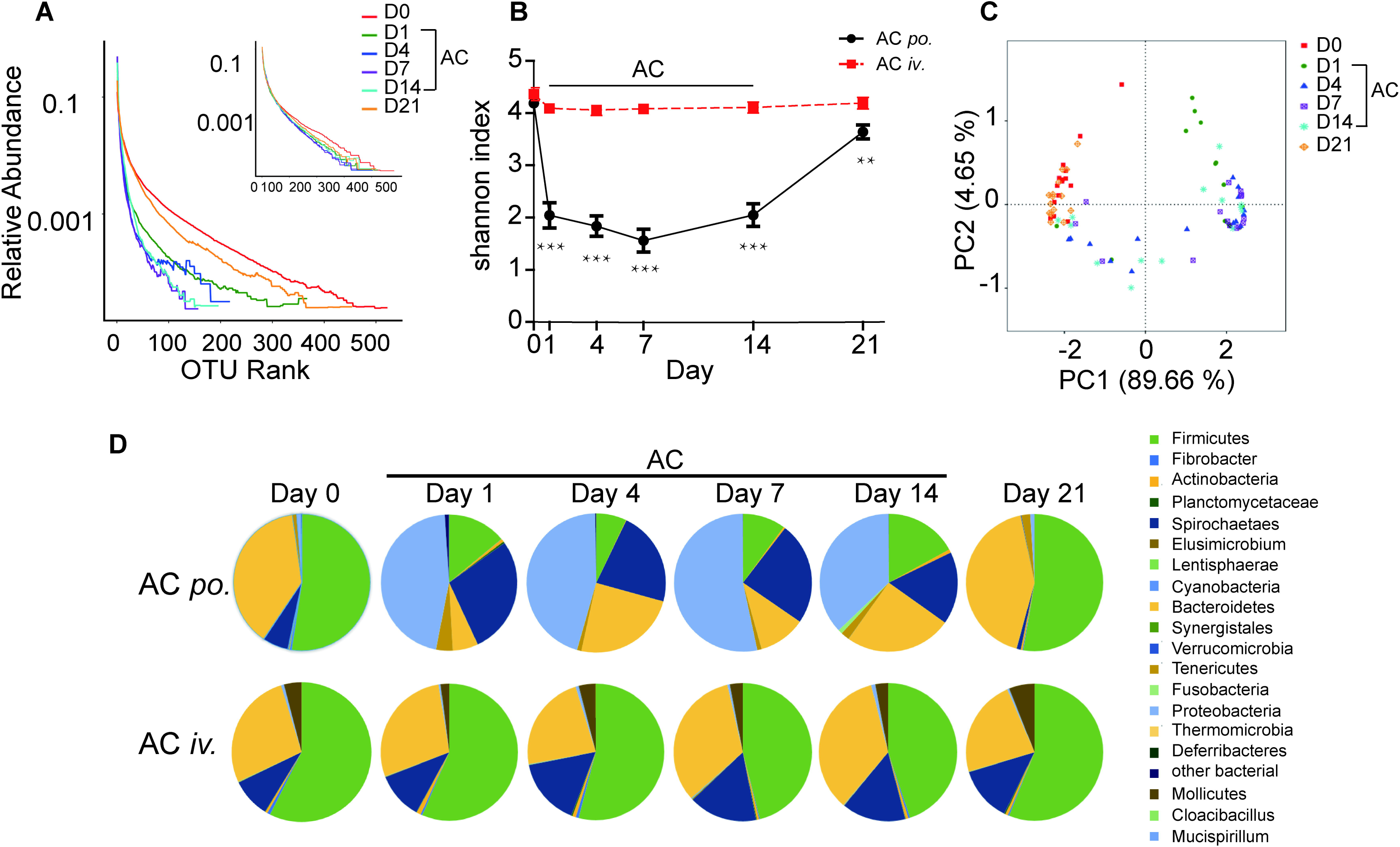
Characteristics of gut microbiota after AC medication. (A) Rank-abundance distribution curve showing the bacterial diversity of AC *po*. group and AC *iv*. group (inserted). The horizon axis represented the OTU abundance in rank order; the vertical axis represented the relative abundance of sequences on a logarithmic scale. (B) Temporal changes of Shannon index after AC medication in AC *po*. group (black line) and AC *iv*. group (red line), **p < 0.01, ***p < 0.0001 compared with day 0. (C) PCoA of the weighted uniFrac β- diversity distance for AC *po*. group. (D)Temporal changes of phyla composition after AC medication. Pie charts showing the phyla found in AC *po*. group and AC *iv*. group. All data were expressed as mean ± SEM, AC *po*. group, n = 15; AC *iv*. group, n = 6 (OTU, operational taxonomic unit; PCoA: principal co-ordinates analysis)

PCoA analysis showed 89.66% and 4.65% of the total variance found could be explained by the first and second axis of the PCoA scatterplots, respectively. Microbiomes at day 21 and day 0 were clustered together, while a clear separation were observed between the microbiota during dosing period and non-dosing period, which indicated oral AC medication induced a significantly reversible microbiol composition change. (Fig. 1C).

The composition of microbe in phylum level was shown in Fig. 1D. At day 0, Firmicutes (51.71 ± 2.66%) and Bacteroidetes (39.25 ± 3.40%) were dominant members. In *po.* group, Firmicutes and Bacteroidetes decreased to 7.14 ± 2.82% (day 4), 8.09 ± 4.98% (day 1) respectively, while Proteobacteria phylum and Spirochaetae phylum increased from 2.48±1.04% (day 0) to 49.79±7.15 % (day 7), and 5.31±1.16% (day 0) to 28.70 ± 6.25 (day 1), respectively during dosing period. Seven days after AC withdrawal, the abundance of Firmicutes, Bacteroidetes, Proteobacteria and Spirochaetae mostly returned to their baseline levels. However, little change was found in *iv.* group during the whole experiment process.

### Oral antibiotics led to increase of BBB permeability

Qualified CSF samples were successfully collected from 12 out of the 15 monkeys (Table 1). AR increase fold elevated significantly to 1.26 ± 0.11 at day 1, and reached its peak of 1.44 ± 0.10 at day 4 in AC *po*. group (p = 0.030 compared to day 0). Thereafter, the level deceased gradually but remained relatively high compared with the baseline level during the remaining dosing period. Seven days after cessation of AC treatment, the fold increase of AR decreased to 1.06 ± 0.08 which approximately back to pre-antibiotics level (Fig. 2A, black curve). In AC *iv*. group, there was just a mild fluctuation of AR with no significant change (Fig. 2A, red curve). In AC *po*. group, AR increased in every monkey during the dosing period, although the change pattern varied greatly among the animals (Suppl. Fig 2). Therefore, AR of pre-antibiotics value (day 0), maximal value, post-antibiotics value (day 21) and the corresponding Shannon index of each monkey were chosen for further correlation analysis. The Shannon index, as well as sobs and chao index, was significantly negatively correlated with the fold increase of AR (r = −0.575, p = 0.004; r = −0.587, p = 0.003; r = −0.531, p = 0.002, respectively Fig.2B-D).

**Fig.2.**
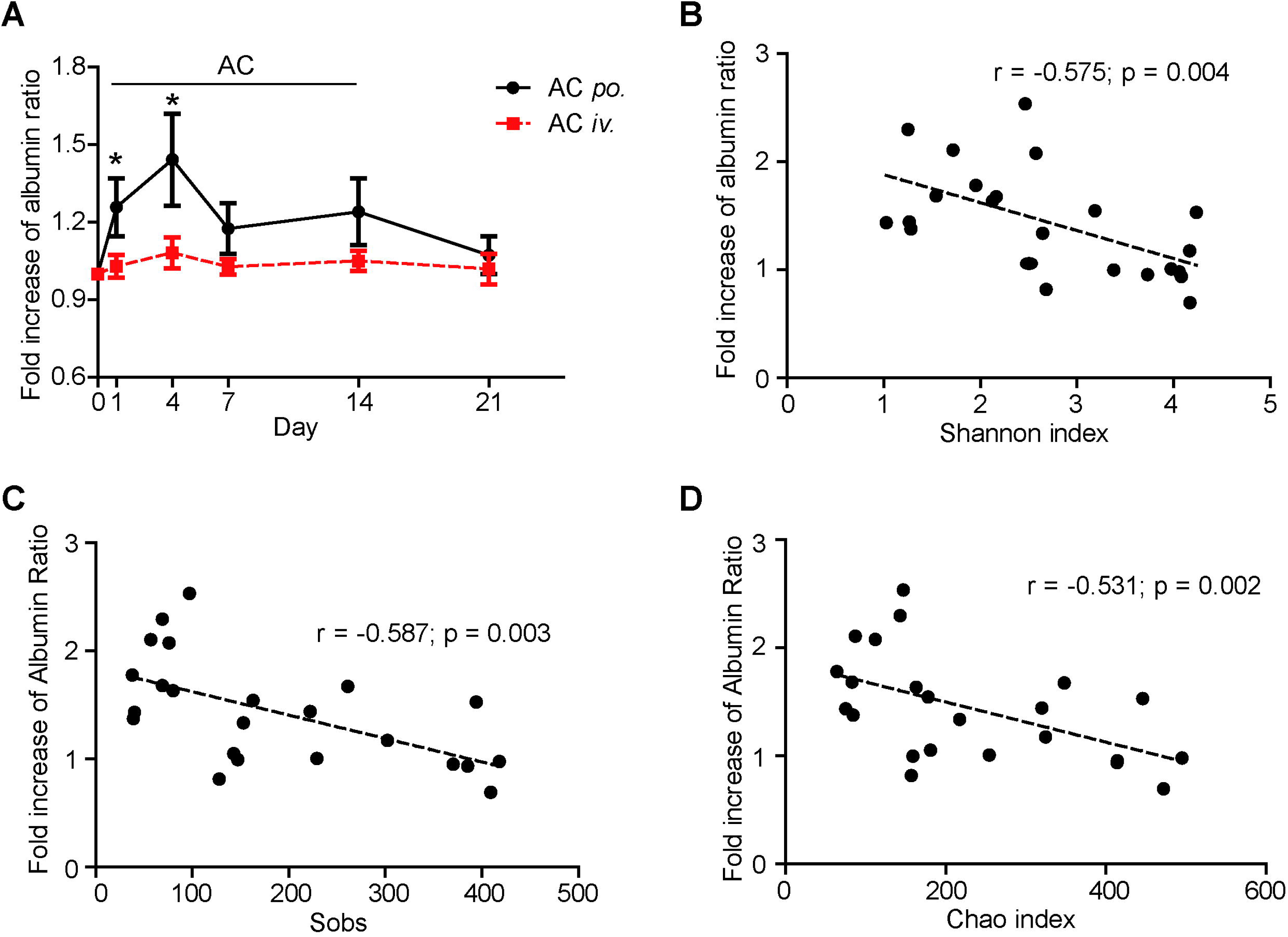
Reversible increase of BBB permeability determined by AR after AC medication. (A) Temporal change of BBB permeability determined by AR in AC *po*. group (black line) and AC *iv*. group (red line), *p < 0.05 compared with day 0 in AC *po*. group. All data were expressed as mean ± SEM, AC *po.* group, n = 12; AC *iv.* group, n = 6 (B-D) The correlations between fold increase of AR and Shannon index, as well as Sobs and Chao index in AC *po*. group.

### BBB permeability increased most significantly at thalamus

After AC administration, a general upward tendency of K_trans_ was observed in all five ROIs at different time points compared with that of day 0 (Fig. 3), although the changing trends were completely discordant among the ROIs (Suppl. Fig 3). There were no significant differences found in the K_trans_ values of caudate nucleus, white matter, and hippocampus, although there was a consistent uptrend change in hippocampal area (p = 0.145, 0.179, 0.106, 0.060 at day 4, day 7, day 14 and day 21 respectively compared with day 0) throughout the experiment (Fig. 3C-E). Notably, in thalamus, the K_trans_ value increased consistently and significantly during dosing period (p = 0.030, 0.003, 0.006 at day 4, day 7, and day 14 respectively compared with day 0) and this phenomenon was also observed at day 21 after cessation of the AC treatment (p = 0.022, Fig. 3F). In addition, there was a statistically significant difference of the K_trans_ value of frontal cortex between day 7 and day 0 (p = 0.013, Fig. 3G).

**Fig. 3.**
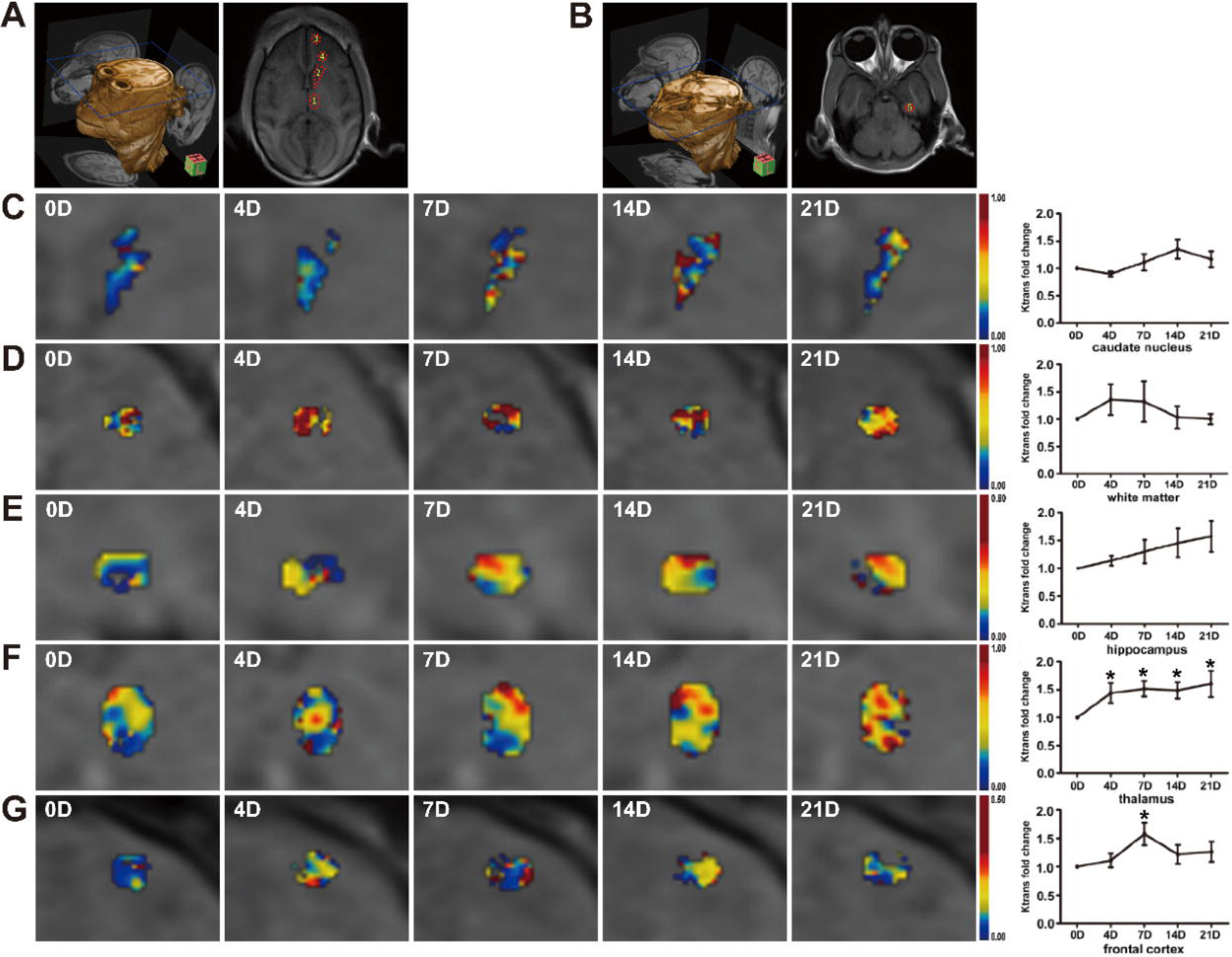
BBB breakdown investigated by DCE-MRI after AC medication. A, B showed representative images of five ROIs encircled (1: thalamus; 2: caudate nucleus; 3: frontal cortex; 4: white matter; 5: hippocampus). C-G: Color map of K_trans_ in regions of interest (ROI), *p < 0.05 compared to day 0, All data were expressed as mean ± SEM, n = 8.

### Oral antibiotic treatment altered microbial metabolism

As SCFAs were shown to be a probable regulators of BBB integrity in germ-free mice^15^, level of the acetic acid, propionic acid and butyrate decreased significantly during dosing period (for acetic acid, p = 0.018, 0.030, 0.011 at day 1, day 4 and day 7 respectively compared with day 0; for propionic acid, p = 0.050, 0.026, 0.002, 0.010 at day 1, day 4, day 7 and day 14 respectively compared with day 0; for butyrate, p = 0.007, 0.022, 0.012 at day 1, day 4 and day 7 compared with day 0) and returned to baseline at day 21 after cessation of the AC (Fig. 4A). Trends in the relative abundance of nine SCFA-producing genera (*Phascolarctobacterium, Subdoligranulum, Faecalibacterium, Blautia, Roseburia, Ruminococcus, Coprococcus, Dorea*, and *Anaerostipes*) were broadly consistent with the trend in the content of SCFAs (Suppl Fig. 4). Positive correlations were observed between three types of SCFAs (acetic acid, propionic acid and butyrate) concentration and relative abundance of the nine genera (r = 0.426, p = 0.001; r = 0.376, p = 0.005; r = 0.327, p = 0.030 respectively, Fig. 4B-D). Considering the above-mentioned nine genera were all from phylum Firmicutes, we analyzed the changes of relative abundance of phylum Firmicutes. On the whole, the trend in relative abundance of phylum Firmicutes was similar with that of acetic acid, propionate acid and butyrate concentration (Fig. 4E) and negative correlation was found between relative abundance of phylum Firmicutes and AR (Fig.4F). Positive correlation was found between acetic acid, propionic acid, butyrate and phylum Firmicutes (r = 0.673, p < 0.001; r = 0.556, p < 0.001; r = 0.708, p < 0.001, Fig. 4G-I). Moreover, the concentration of acetic acid and propionic acid was significantly negatively correlated to the fold increase of AR (r = −0.748, p = 0.005 and r = −0.681, p = 0.015, respectively, Fig. 4J, K), while no significantly negative correlation was found between butyrate concentration and AR (Fig. 4L). Besides, we also evaluated the correlations between phylum Firmicutes, acetic acid, propionate acid, butyrate and the fold increase of K_trans_ (hippocampus region and thalamus region). Negative correlation was found between relative abundance of phylum Firmicutes, acetic acid, propionic acid and the fold increase of K_trans_ in the thalamus region, while only propionic acid was negatively correlated to the fold increase of K_trans_ in the hippocampus region (Supplementary table).

**Fig. 4.**
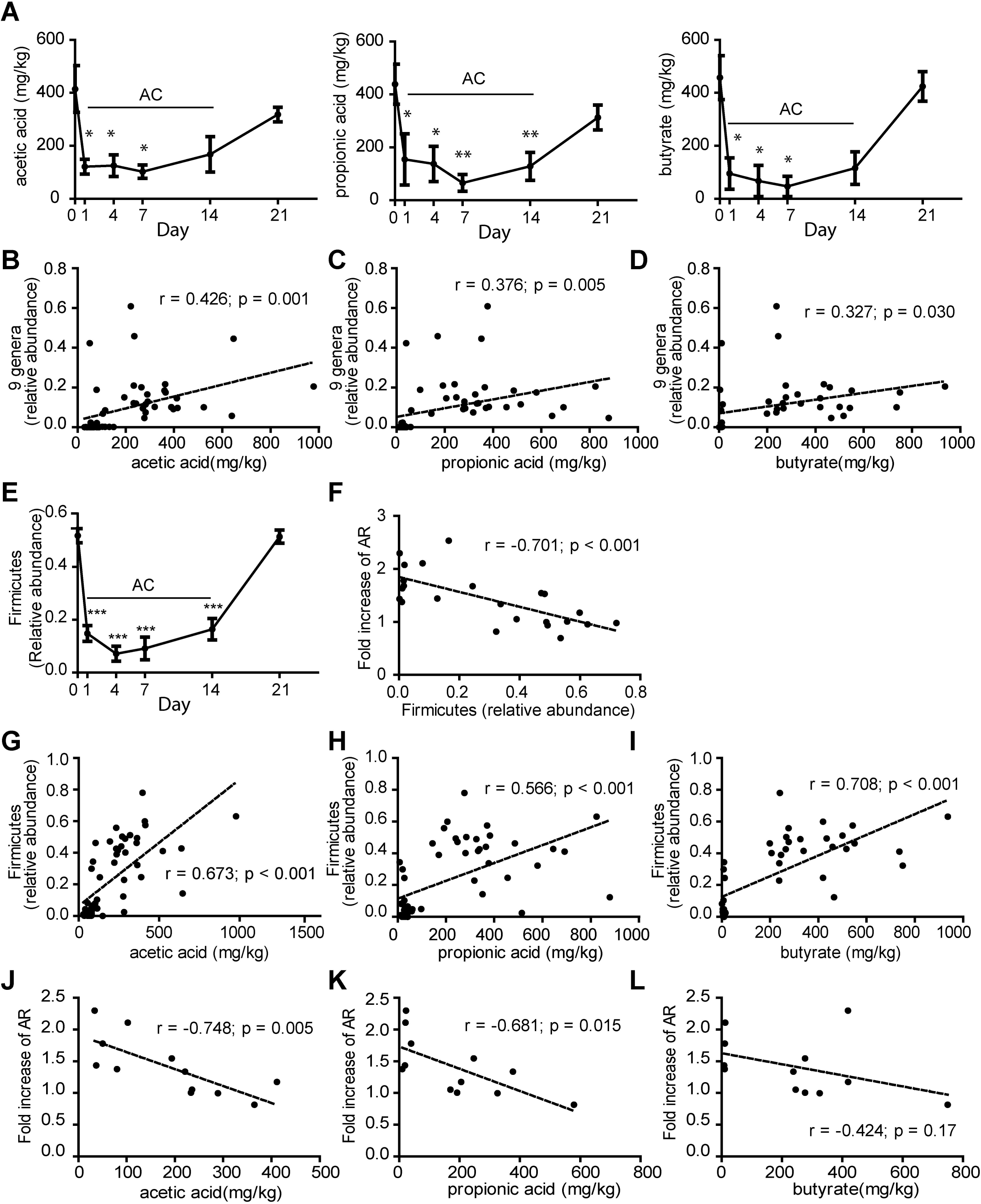
Changes of acetic acid, propionate acid, butyrate and its correlation with genera, phylum Firmicutes and AR. (A) Temporal changes of acetic acid, propionic acid and butyrate concentration, *p < 0.05, **p < 0.01 compared with day 0, n = 9. (B-D) Correlation between acetic acid, propionic acid, butyrate concentration and relative abundance of 9 genera producing SCFAs, n = 9. (E) Changes of relative abundance of phylum Firmicutes, ***p < 0.001 compared with day 0, n = 15. (F) Correlation between phylum Firmicutes and AR, n = 12. (G-I) Correlations between relative abundance of phylum Firmicutes and acetic acid, propionate acid, butyrate, n = 9. (J-L) Correlation between acetic acid, propionate acid, butyrate and fold increase of AR, n = 6.

## DISCUSSION

Effect of gut microbiome on BBB has been controversial, with evidence showing abnormal BBB development and decreased level of *occludin* and *claudin-5* mRNA in amygdala and hippocampus germ-free mice, while upregulated proteins Occludin and Claudin-5 were seen in mice receiving low-dose oral penicillin in early life^11, 12^. To the best of our knowledge, this is the very first study to investigate on direct evidence of effects of gut microbiol alterations on BBB permeability and thus CNS.

In the current study, BBB permeability was determined using a standard method involving measuring AR, which was similar and comparable to the method used in current clinical practice. Although some techniques such as microdialysis could be used for rodent CSF sampling, it is only suitable for analysis of metal ions, neurotransmitters, and other small molecules, rather than protein analysis. Non-human primates are not only more translational relevant, it can provide as much as 800 μL CSF at each sampling time point by using the minimally invasive catheterization method developed in our lab^9^, making it possible to measure AR and thus BBB permeability directly.

A drastic shift of the biodiversity and richness of gut microbiome was induced by oral AC treatment at a comparable dosage. Accordingly, BBB permeability determined by continuous AR and K_trans_ increased significantly in oral, but not intravenous, AC treated animals. These findings suggested that antibiotics-induced gut microbiol composition change not only could affect CNS and BBB permeability, but also could affect them directly rather than through contact between blood and brain endothelium. Meanwhile, a healthy gut microbiome baseline and a reversion trend of BBB permeability change towards normal status after medication discontinuation was observed in healthy adult monkeys, indicating that BBB was under relatively normal physiological conditions before antibiotic treatment. These results might suggest a totally different BBB permeability disruption mechanism in adults, which might be more correlated with a direct gut microbiol composition alterations induced by antibiotics oral administration, comparing to BBB dysplasia before adulthood as previously reported^11^.

Antibiotics have been a pillar of modern medicine as a means to treat bacterial infections in humans and animals and have been widely used on an enormous scale around the world for almost a century^16^. Prophylactic treatment, including after stroke, has been one of the most frequent usage scenarios of antibiotics. As we previously reported, the extent of BBB permeability disruption was highly correlated to ischemic outcome in monkeys subjected to ischemic stroke^9^. Considering that only orally administrated AC interfered with gut microbiome and thus BBB permeability while intravenous AC achieving the same blood concentration did not, as presented in our results, it could be reasonable to speculate that intravenous prophylactic antibiotics might be associated with better patient outcomes by preserving relatively intact BBB function. Meanwhile, the modification of microbiota caused by oral antibiotics was partially reversible after medication discontinuation in current study, which was consistent with the recent study of MacPherson et al. using AC treatment in healthy adults^17^. And it has been established that antibiotic characteristics including, but not limited to, class, pharmacokinetics, pharmacodynamics and range of action, as well as their dosage, duration and administration route, affect their final effects on gut flora^18^ and thus BBB permeability. It is not a surprise to see the lack of rapid reversion when other antibacterial agents (cefprozil, clarithromycin, metronidazole, ciprofloxacin etc.) and dosing scheme, were used^19-21^. Therefore, it might also be important to consider possible impact on gut microbiome when choosing antibiotics, thus to avoid BBB permeability interference as possible.

A persistent worsening of gut microbiol composition alterations and BBB permeability during the whole AC administration period was noted, indicating an accumulation of gut microbiome composition and BBB function compromise throughout antibiotics treatment. As short course of antibiotics usage might be able to gain curative effect comparable to longer course^22^, shorter duration of antibiotics treatment might should be preferred in order to protect original gut microbiome and preserve normal BBB permeability.

In current study, the BBB permeability change did not happen homogenously in the whole brain. Although a general upward tendency of K_trans_ was observed in all five ROIs (thalamus, caudate nucleus, frontal cortex, white matter, hippocampus), the pattern differed greatly in different regions. Thalamus, an area critical for relaying sensory and motor signals to cortex, regulation of consciousness and alertness, showed the most statistically significant increase of K_trans_. Hippocampus, a region involved in learning and memory showed a very consistent increase trend although differed a lot in each single animal. It is noteworthy that the hippocampal BBB breakdown happens in AD as well^23^. This observation might explain why disruptions of gut microbiome are strongly correlated with neurological and psychological disease such as AD, ASD, and depression etc^24^. In mid-aged or senior people, where a more frequent use of oral antibiotics happens, this gut microbiol alteration-induced BBB permeability increase might be an important factor contributing to multiple pathophysiological processes of neurodegenerative diseases, and might serve as a future therapeutic target.

Circulating metabolites may explain the “bottom-up” communication. For instance, Bercik’s team^25^ has demonstrated that changes of brain chemistry induced by alteration of the gut microbiome could occur independently of vagal or sympathetic neural pathways and in the absence of any immune response, strongly suggesting a potential mechanism by which microbiome influence the BBB is by producing metabolites that can alter CNS function. SCFAs, such as butyrate, acetate, and propionate, produced through the fermentation of dietary fibers by the gut microbiota have been shown to improve BBB integrity in germ-free mice^15^. A recent report has shown the protective effects of propionate upon BBB endothelial cells against LPS-induced barrier disruption via a CD14-dependent mechanism^26^. In current study, SCFAs levels decreased significantly after AC treatment. This may result from the reduction in some SCFA-producing gut community members of SCFA-producing members. Besides, acetic acid and propionate acid were found to be correlated with AR while butyrate was not, which was different from germ-free mice^14^.

## CONCLUSIONS

In conclusion, our study provides strong evidence that antibiotics-induced gut microbiol composition change, especially the decreasing of acetic and propionic acid producing phyla and genera, leads to increase of BBB permeability, which may be an important component in some of the emerging links between intestinal microbiol composition alterations and pathologies as significant as depression, AD, PD, and ASD. Therefore, the modulation of gut microbiota should be considered a new, striking therapeutic avenue to be used not only for infectious diseases, but also for neurological disorders associated with the disruption of BBB. Moreover, a wiser use of antibiotics in clinical practice, especially in BBB vulnerable young children, and patients with known pathological BBB disruptions, is advocated and need special caution. which may contribute to a variety of neurological and psychological diseases.

## Abbreviations

BBB: Blood-brain barrier;
AC: amoxicillin-clavulanic acid;
AR: CSF/serum albumin ratio;
K_trans_: volume transfer constant;
CNS: central nervous system;
AD: Alzheimer’s disease;
PD: Parkinson’s disease;
ASD: autism spectrum disorder;
GF: germ-free;
CSF: cerebrospinal fluid;
ROIs: regions of interest;
SCFAs: short chain fatty acids.

## Acknowledgements

The authors are grateful to Jinge Zhang, Jie Zhang and Huaiqiang Sun for their valuable assistance in conducting experiments.

## Availability of data and materials

All data generated or analyzed during this study are included in this published article and its supplementary information files.

## Authors’ contributions

QW helped to design the research, coordinated all experiment operations, conducted and performed all statistical analysis, and wrote the manuscript; YQZ helped to design the research, assisted in conducting experiments, collecting data and analyzing samples, and helped to prepare the manuscript; YBZ assisted in performing the animal management and CSF sampling preparation; CCX conducted MRI scanning; QL was responsible for catheterization procedures; ZQD and WHK helped on medication design; CY and DS helped on sample analysis; HXL helped on animal medication and welfare; ZHZ designed the entire study, supervised all segments of the study, and revised the manuscript.

## Ethics approval

Animal use followed the NIH Guide for the Care and Use of Laboratory Animals. All experimental protocols were approved by the Institutional Animal Care and Use Committee (IACUC), Animal Experiment Center of Sichuan University and met institutional and national guidelines.

## Consent for publication

All authors have read the manuscript and indicated consent for publication.

## REFERENCE

1. Schmidt, C. 2015. Mental health: thinking from the gut. Nature. 518: S12.

2. Smith, P. A. 2015. The tantalizing links between gut microbes and the brain. Nature. 526: 312–314.

3. Chesnokova, V. & R. N. Pechnick. 2016. New Signaling Pathway for Gut–Brain Interactions. Neuropsychopharmacology. 41: 372.

4. Cryan, J. F. & T. G. Dinan. 2015. More than a gut feeling: the microbiota regulates neurodevelopment and behavior. Neuropsychopharmacology. 40: 241–242.

5. Cryan, J. F. & T. G. Dinan. 2012. Mind-altering microorganisms: the impact of the gut microbiota on brain and behaviour. Nature reviews. Neuroscience. 13: 701–712.

6. Obermeier, B., R. Daneman & R. M. Ransohoff. 2013. Development, maintenance and disruption of the blood-brain barrier. Nature medicine. 19: 1584–1596.

7. Rosenberg, G. A. 2012. Neurological diseases in relation to the blood-brain barrier. Journal of Cerebral Blood Flow & Metabolism Official Journal of the International Society of Cerebral Blood Flow & Metabolism. 32: 1139.

8. Zhong, Z., et al. 2009. Activated protein C therapy slows ALS-like disease in mice by transcriptionally inhibiting SOD1 in motor neurons and microglia cells. Journal of Clinical Investigation. 119: 3437–3449.

9. Zhang, Y., et al. 2017. Temporal analysis of blood-brain barrier disruption and cerebrospinal fluid matrix metalloproteinases in rhesus monkeys subjected to transient ischemic stroke. J Cereb Blood Flow Metab. 37: 2963–2974.

10. Heye, A. K., et al. 2014. Assessment of blood-brain barrier disruption using dynamic contrast-enhanced MRI. A systematic review. NeuroImage. Clinical. 6: 262–274.

11. Frohlich, E. E., et al. 2016. Cognitive impairment by antibiotic-induced gut dysbiosis: Analysis of gut microbiota-brain communication. Brain, behavior, and immunity. 56: 140–155.

12. Leclercq, S., et al. 2017. Low-dose penicillin in early life induces long-term changes in murine gut microbiota, brain cytokines and behavior. Nature communications. 8: 15062.

13. Jernberg, C., et al. 2007. Long-term ecological impacts of antibiotic administration on the human intestinal microbiota. The ISME journal. 1: 56–66.

14. Braniste, V., et al. 2014. The gut microbiota influences blood-brain barrier permeability in mice. Sci Transl Med. 6: 263ra158.

15. Volpe, B. T. 2014. The Gut Microbiota and the Blood-Brain Barrier. Science Translational Medicine. 6.

16. Blaser, M. J. 2016. Antibiotic use and its consequences for the normal microbiome. Science. 352: 544.

17. MacPherson, C. W., et al. 2018. Gut Bacterial Microbiota and its Resistome Rapidly Recover to Basal State Levels after Short-term Amoxicillin-Clavulanic Acid Treatment in Healthy Adults. Scientific reports. 8: 11192.

18. Ianiro, G., H. Tilg & A. Gasbarrini. 2016. Antibiotics as deep modulators of gut microbiota: between good and evil. Gut. 65: 1906.

19. Pérezcobas, A. E., et al. 2013. Differential Effects of Antibiotic Therapy on the Structure and Function of Human Gut Microbiota. Plos One. 8: e80201.

20. Raymond, F., et al. 2016. The initial state of the human gut microbiome determines its reshaping by antibiotics. Isme Journal. 10: 707.

21. Willing, B. P., S. L. Russell & B. B. Finlay. 2011. Shifting the balance: antibiotic effects on host-microbiota mutualism. Nature Reviews Microbiology. 9: 233–243.

22. Dawson-Hahn, E. E., et al. 2017. Short-course versus long-course oral antibiotic treatment for infections treated in outpatient settings: a review of systematic reviews. Fam Pract. 34: 511–519.

23. Montagne, A., et al. 2015. Blood-brain barrier breakdown in the aging human hippocampus. Neuron. 85: 296–302.

24. Sharon, G., et al. 2016. The Central Nervous System and the Gut Microbiome. Cell. 167: 915–932.

25. Bercik, P., et al. 2011. The intestinal microbiota affect central levels of brain-derived neurotropic factor and behavior in mice. Gastroenterology. 141: 599-609, 609 e591-593.

26. Hoyles, L., et al. 2018. Microbiome–host systems interactions: protective effects of propionate upon the blood–brain barrier. Microbiome. 6: 55.

