## Supplementary material for "Antibiotics-induced alterations in gut microbiol composition leads to blood-brain barrier permeability increase in rhesus monkeys"

**Supplementary methods**

**Serial CSF Sampling from Cisterna Magna**

Briefly, a clinically routine lumbar puncture was operated on monkeys followed by a catheterization under guidance of X-ray. The catheter reached the cisterna magna and its tip kept floating in CSF afterwards. A sampling port was embedded under skin beside the puncture site and connected to the tube, through which the cisterna magna CSF could be extracted according to experiment plan during the whole study (Fig. S1). To assess the dynamic change of BBB permeability, blood and cisterna magna CSF were sampled at day 0, day 1, day 4, day 7, day 14 and day 21.

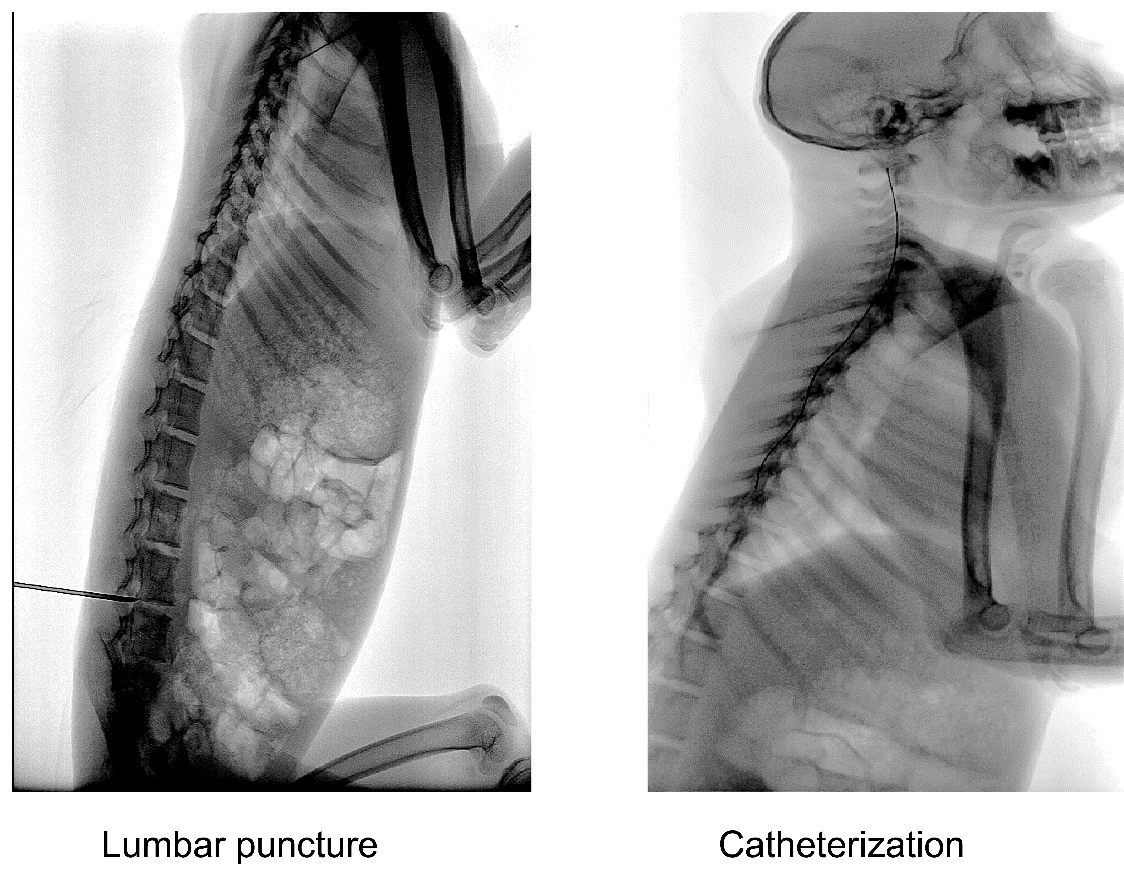

Fig. S1 Seral CSF sampling from cisterna magna

**16S rRNA gene sequence analysis**

The V4-V5 hypervariable regions of the bacteria 16S rRNA gene were amplified by PCR (95 °C for 2 min, followed by 25 cycles at 95 °C for 30 s, 55 °C for 30 s, and 72 °C for 45 s, and a final extension at 72 °C for 10 min) using barcodes primers 515F (5’-GTGCCAGCMGCCGCGGTAA-3’) and 926R (5’-CCGTCAATTCMTTTRAGTTT-3’). PCR reactions were performed in triplicate 20 μL mixture containing 4 μL of 5 × FastPfu Buffer, 2 μL of 2.5 mM dNTPs, 0.8 μL of each primer (5 μM), 0.4 μL of FastPfu Polymerase and 10 ng of template DNA. The resulted PCR products were extracted from a 2 % agarose gel and further purified using the AxyPrep DNA Gel Extraction Kit (Axygen Biosciences, Union City, CA, USA) and quantified using QuantiFluor™-ST (Promega, USA) according to the manufacturer’s protocol.

Sequences of purified amplicons were quality-filtered using the QIIME v1.8 with the following criteria: (i) The reads were truncated at any site receiving an average quality score < 20 over a 50 bp sliding window. (ii) Primers were exactly matched allowing 2 nucleotide mismatching, and reads containing ambiguous bases were removed. (iii) Sequences whose overlap longer than 10 bp were merged according to their overlap sequence. Operational taxonomic units (OTUs) were clustered with 97 % similarity cutoff using UPARSE（version 7.1 http://drive5.com/uparse/) and chimeric sequences were identified and removed using UCHIME. The taxonomy of each 16S rRNA gene sequence was analyzed by RDP Classifier algorithm (http://rdp.cme.msu.edu/) against the Greengenes 16S rRNA database.

**Gas chromatography analysis of SCFAs**

Short-chain fatty acids (SCFAs), including acetic acid, propionate acid and butyrate, in monkey fecal samples were quantified by gas chromatography (GC). A Bruker 430-GC system (Billerica, MA, USA), equipped with a flame ionization detector (FID) and an automatic liquid sampler, was used. A J&W DB-FFAP capillary column (Agilent Technologies Inc., USA; Part # J122-3232, 30 m × 0.25 mm × 0.25 μm film thickness) was used. Two gram of feces which was previously frozen and thawed on ice, was dissolved, vortexed and mixed in 10 mL of deionized water. The resulting slurry was sonicated for 10 min and centrifuged at 12000 rpm for 5 minutes. Two mL diethyl ether and 0.5 mL 50 % sulfuric acid solution was added to 5 mL of collected supernatant and the mixture was vortexed for 1 min prior to centrifugation. After standing for 30 min, the upper layer of ether extract was passed through a 0.45 μm filter prior analysis. The injection volume was 2 μL. Each sample run was preceded with a wash run of 1 % formic acid. The oven temperature was held at 100 °C for 1 min, then increased to 200 °C at 15 °C / min and held for 2 min. The temperature for the FID and the injection port was 280 °C and 270 °C, respectively. The flow rate of helium, hydrogen, and air was 40 mL/min, 40 mL/min, and 400 mL/min, respectively. SCFAs were identified and calculated using external standards consisting of authentic acetate, propionate, and butyrate.

**Supplementary figures**

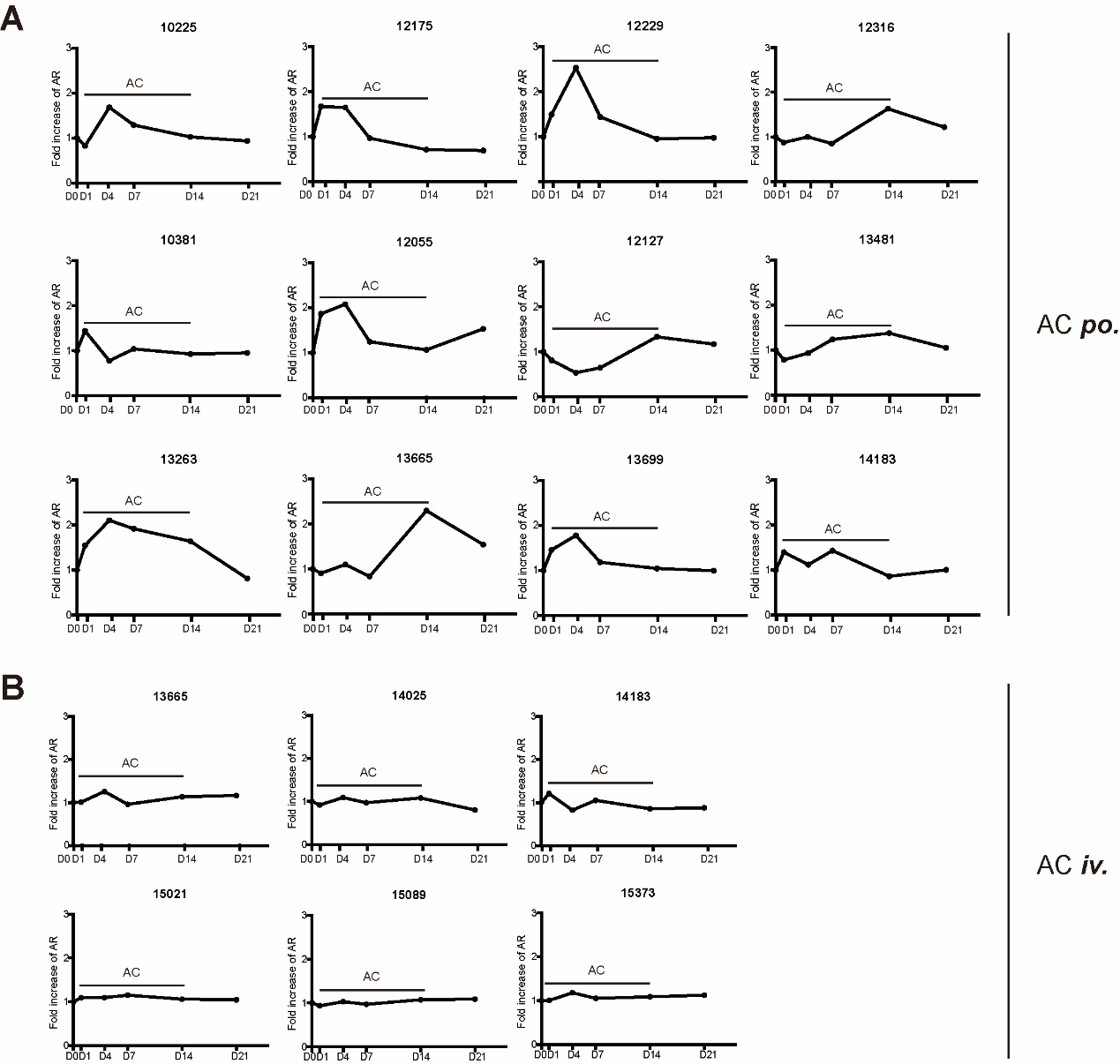

Fig. S2. The fold change of AR of each monkey in AC *po.* group (A) and AC *iv.* group (B)

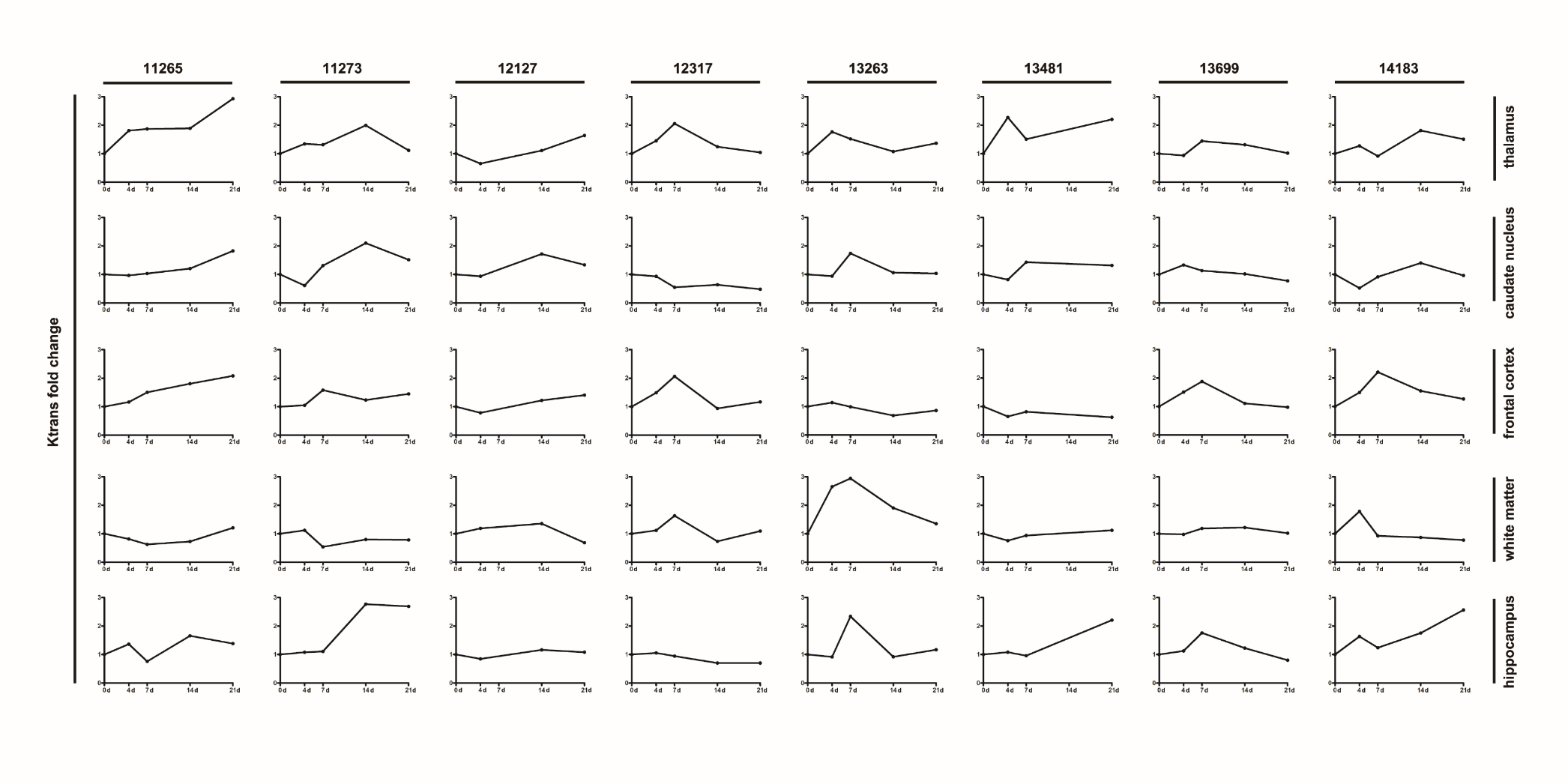

Fig. S3. The fold change of Ktrans in five ROIs of each monkey in AC *po.* group.

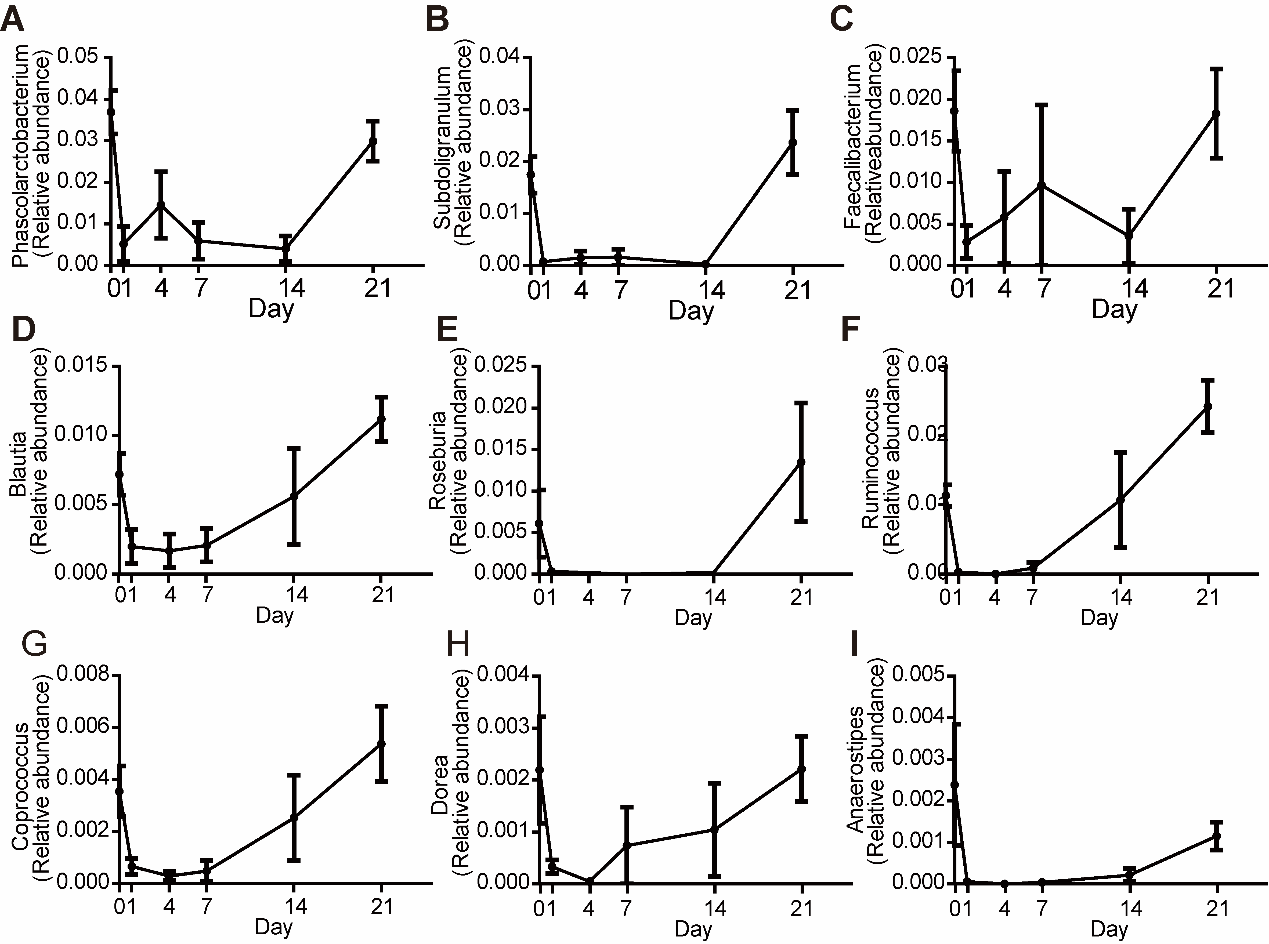

Fig. S4 The temporal change of nine SCFA-producing genera (*Phascolarctobacterium*, *Subdoligranulum*, *Faecalibacterium*, *Blautia*, *Roseburia*, *Ruminococcus*, *Coprococcus*, *Dorea*, and *Anaerostipes*). All data were expressed as mean ± SEM, n = 15

**Supplementary table: The correlations between K_trans_ in hippocampus / thalamus and changes of Firmicutes, acetic acid, propionate acid and butyrate（n=8）.**

| K_trans_ | Firmicutes | | | acetic acid | | | propionic acid | | | butyrate | |
| --- | --- | --- | --- | --- | --- | --- | --- | --- | --- | --- | --- |
|  | r | p | r | | p | r | | p | r | | p |
| hippocampus | -0.229 | 0.282 | -0.35 | | 0.093 | -0.486 | | 0.016 | -0.323 | | 0.143 |
| thalamus | -0.459 | 0.024 | -0.418 | | 0.042 | -0.418 | | 0.042 | -0.365 | | 0.095 |
